# Sociability is a multidimensional trait in *Drosophila melanogaster*

**DOI:** 10.64898/2025.12.18.695080

**Authors:** Tiphaine P.M. Bailly, Sanne J.C. Lamers, Adithya Sarma, Anne C.M. Jansen, Koen Freerks, Michael van Dijk, Rampal S. Etienne, Bregje Wertheim, Jean-Christophe Billeter

## Abstract

Sociability—the propensity of an individual to engage in group activities—is a trait present in all social species. In humans and many animals, sociability varies between individuals yet remains consistent across contexts, qualifying it as a personality trait. Sociability influences health and physiology, but the mechanisms underlying sociability and its inter-individual variation remain poorly understood. The genetically tractable fruit fly, *Drosophila melanogaster*, is increasingly used to study social behavior and exhibits a wide range of sociability phenotypes. However, previous studies have relied on distinct behavioral paradigms, limiting cross-context comparisons and motivating a more extensive characterization of sociability in this species. Here, we quantified sociability in *D. melanogaster* using a multidimensional approach encompassing three paradigms that capture engagement in group activities across contexts: (1) preference for communal versus solitary egg-laying, (2) egg-laying latency in a group, and (3) frequency and duration of spontaneous social interactions and interindividual distance. We assessed these behaviors in 105 lines of the *Drosophila* Genetic Reference Panel and observed substantial variation in responses to conspecific presence across paradigms. Sociability-related behaviors differed between genetically distinct lines, indicating a genetic component. However, the three sociability traits were uncorrelated, demonstrating that sociability in *D. melanogaster* is multidimensional. These findings suggest that sociability is not governed by a single central mechanism, but instead arises from multiple context-dependent pathways.

## Introduction

Group formation, or aggregation, occurs in most organisms from bacteria to humans, making gregariousness one of the most ubiquitous patterns in life (M. Chen & Sokolowski, 2022; Frank, 2007; Parrish & Edelstein-Keshet, 1999; West et al., 2007). Although individuals in groups must compete for food, space and mates, and have higher risk of infectious diseases (Clutton-Brock & Huchard, 2013; Clutton-Brock & Scott, 1991; Otterstatter & Thomson, 2007; Stockley & Bro-Jørgensen, 2011), individuals benefit from associating with others for several reasons including reducing predation risk, removing ectoparasites, enhancing foraging and reproductive success (Allee, 1938; Danchin et al., 2004; Krause & Ruxton, 2002; Sakata & Katayama, 2001; Wertheim, 2001). The balance of these costs and benefits is likely at the root of the diversity in sociality, the degree to which individuals in a species associate in social groups and form cooperative societies. This ranges from mostly solitary species, whose social interactions are limited to mating and aggression, to eusocial species, who spend their whole life in groups and cannot survive alone.

In addition to the diversity of sociality across species, there is within-species variation in the propensity of individuals to associate with other individuals outside of mating and aggressive encounters: termed sociability (Gartland et al., 2022). Individuals displaying a strong tendency to interact with conspecifics are deemed highly sociable, while others prefer to stay away from groups. In humans, sociability is defined as a tendency to affiliate with others and to prefer being with others over remaining alone (Boswell et al., 2020; Cheek & Buss, 1981). Human sociability is measured by both the quality and the quantity of social contact, which is mostly measured by subjective measures (Boswell et al., 2020; Bralten et al., 2021). For example, a genome wide association study aimed at mapping the genetic basis of sociability quantified sociability through four questions regarding the frequency of friend/family visits, the number and type of social venues visited, response to social embarrassment, and perceived loneliness (Bralten et al., 2021). This illustrates that sociability is a complex behavioral conglomerate. In non-human animals, sociability measures are divided into two subgroups (Gartland et al., 2022). The first include social approach assays, in which a focal individual is placed in an environment with one or more conspecific and sociability is quantified by the distance between them. The second subgroup consists of social network and collective behavior analyses, where social interactions among multiple individuals are tracked over time, and individual sociability scores are calculated from repeated observations with the same or different groupmates. There is a wide range of methodologies to quantify sociability across animal behavior research, but it is unclear whether these sociability measures correlate with each other and/or measure the same dimension of sociability.

One way to better define sociability may be to identify what molecular and cellular mechanisms are causal to its variation through identification of genes influencing sociability. Genetic variation plays an important role in sociability and several components of this trait are heritable in humans and animals. For instance, loneliness, the propensity to engage in social interactions and prosocial behaviors (*e.g.* helping, sharing, donating, co-operating and volunteering) are heritable and can be observed among members of the same family (Abdellaoui et al., 2018; Boomsma et al., 2005; Distel et al., 2010; Ebstein et al., 2010; Fowler et al., 2009; Goossens et al., 2015; Ordoñana et al., 2013; van den Berg et al., 2016). In keeping with the high heritability of sociability, genes that influence sociability have been identified. In vertebrates such as voles, chimpanzees or humans, the genetic pathways of the neuropeptide oxytocin and neurohormone vasopressin vary between individuals and explain differences in social behavior among individuals of the same species, and are implicated in the regulation of sociability (Bakermans-Kranenburg & van Ijzendoorn, 2014; Caldwell, 2012; Donaldson & Young, 2008; Staes et al., 2015). In *Caenorhabditis elegans,* sociability varies between individuals and populations, and solitary *vs* social foraging strategies are determined by natural variation in the *Neuropeptide-y* gene (de Bono & Bargmann, 1998). Sociable phenotypes are observed in the fruit fly, *Drosophila melanogaster,* and fruit fly’s genotypes naturally differ in their degree of sociability (Anderson et al., 2017; Saltz, 2011; Scott et al., 2018; Torabi-Marashi et al., 2025; Wice & Saltz, 2021). Sociability levels vary significantly between genetically distinct fruit fly lines, with broad-sense heritability estimates of 0.24 for males and 0.21 for females (Scott et al., 2018). To determine sociability, studies have used social network analyses (Pasquaretta et al., 2016; Rooke et al., 2024; Schneider et al., 2012; Wice & Saltz, 2021), measured inter-fly distance (Anderson et al., 2017; Fernandez et al., 2017; Jiang et al., 2020; Simon et al., 2012; Sun et al., 2020), examined how aggressive behaviors influenced fly distributions among food patches(Saltz, 2011), studied how social environment influenced food search behavior (Lihoreau et al., 2016; Tinette et al., 2004) or focused on joining others or be alone at a food patch (Scott et al., 2018; Torabi-Marashi et al., 2025). Although all of the *Drosophila* studies mentioned above highlighted variation in sociability, they each quantified sociability through different and unidimensional assays. It is therefore currently unknown whether different “sociability” measures correlate, and hence, whether sociability can be considered a unidimensional trait in *D. melanogaster*.

In this study, we investigated whether variation in sociability in *D. melanogaster* generalizes over three paradigms relying on different sensory modalities that each quantify sociability in a different context. We quantified communal egg-laying preference by exposing a mated female to a choice to lay eggs near aggregation pheromones or away from them, a behavior relying on olfactory detection of volatile pheromones (Duménil et al., 2016; Verschut et al., 2023a). Laying eggs with others is a form of cooperation because it allows the resulting offspring to cooperate in fending off fungus and for burrowing through food (Dombrovski et al., 2017, 2020; Trienens et al., 2017; Wertheim et al., 2002). In parallel, we quantified latency to lay eggs in a group of flies, a behavior in which females lay eggs faster in groups than alone during the day, relying on visual cues for both the detection of other flies and of diurnal cues (Bailly et al., 2023). Finally, we used automated behavioral tracking of 4 individuals to quantify their number and duration of spontaneous social interactions as well as interindividual distance and nearest-neighbor distance. We performed these analyses on strains of the *Drosophila* Genetic Reference Panel (DGRP), which are inbred, sequenced lines that have been used by many laboratories to map the genetic basis of a wide range of phenotypes in different environments (Huang et al., 2014, 2020; Mackay et al., 2012). These strains capture the natural genetic variation found in their original North American population and allow to repeatedly quantify sociability in different paradigms for each of the represented genotypes. Here, we report that three different paradigms reveal extensive interline sociability variation among DGRP lines, but that the three sociability measures do not correlate with each other, suggesting that sociability is a multidimensional trait.

## Materials and methods

### *Drosophila* rearing and stocks

Flies were reared on food medium (agar 10g/L, sucrose 15g/L, glucose 30g/L, cornmeal 15g/L, wheat germ 10g/L, dried yeast 35g/L, soy flour 10g/L, molasses 30g/L, propionic acid 5mM and Tegosept 2g/L) in a 12:12h light/dark cycle at 25°C. Virgin adults were collected using CO2 anaesthesia and aged in same-sex groups of 20 in food vials (25×95 mm) containing food medium (23×15 mm) for 5 to 7 days before testing. Flies used in this study were from the wild-type *Oregon-R* (*OR*), *Canton-S* (*CS*), *w^1118^*, *w^1118^*; *OR* (*w^1118^* backcrossed 10 times to *OR*) lab strains, or from the *Drosophila* Genetic Reference Panel (DGRP). We report data on 105 out of the 205 available DGRP lines (see Table S1). Lines with reduced activity or infertility were excluded, as these traits prevented reliable behavioral quantification. DGRP females used in the different assays were all mated with males of *OR*, to prevent possible bias induced by males during mating. Sociability of these 105 DGRP lines was quantified through three assays that are presented and described below: communal egg-laying preference, communal egg-laying latency and spontaneous social interactions. To mitigate an effect of day and time of day, trials were randomly spread out over multiple days with a maximum of two or five replicates per DGRP line per day depending on the paradigm.

### Sociability assay 1: Communal egg-laying preference

Six DGRP virgin females were mated with 6 *OR* males for 1 hour at 25°C in a 55×14 mm Petri dish containing a 23×1 mm circular fly food patch. Mating was confirmed by checking for a mating plug under a UV flashlight (Lung & Wolfner, 2001). Afterwards, each female was isolated for 24 hours at 25°C (12:12 light:dark cycle) in a 1 ml screw cap tube with a yeast-extract paste (in 3.5% agar) on the lid to prevent egg-laying. Each mated female was then transferred to a 57×38×17 mm rectangular egg-laying dish containing two black fly food patches (black food: 20 g/L activated charcoal, with cornmeal and wheat germ omitted) separated by a 3.5% agar patch (Figure 1A; Verschut et al. 2023). One food patch was enriched with mated-female pheromone extract, while the other received control hexane. To prepare the pheromone extract (Verschut et al., 2022), 6-day-old mated *OR* females (500–900 individuals) were submerged in hexane (12 µL per fly) in a 20 ml glass bottle and vortexed for 2 minutes. The hexane extract containing cuticular hydrocarbons was then aliquoted into 1 ml glass vials, and stored at –20°C. For each assay, 10 µL of pheromone extract and 10 µL of control hexane were applied to separate 5.5 mm circular filter papers (with a parafilm backing), and the hexane was evaporated under a nitrogen flow. The assay dish was placed in a chamber (63 cm deep × 71 cm high × 120 cm long) illuminated with white and red (>620 nm) LEDs on a 12:12 light:dark cycle at 25°C with continuous airflow to prevent odor accumulation. The chamber was cleaned with ethanol after each experiment. Egg-laying preference was determined by manually counting the eggs on each patch and calculating a preference index as: ((# eggs on pheromone patch) – (# eggs on control patch))/{total # eggs}.

**Figure 1:**
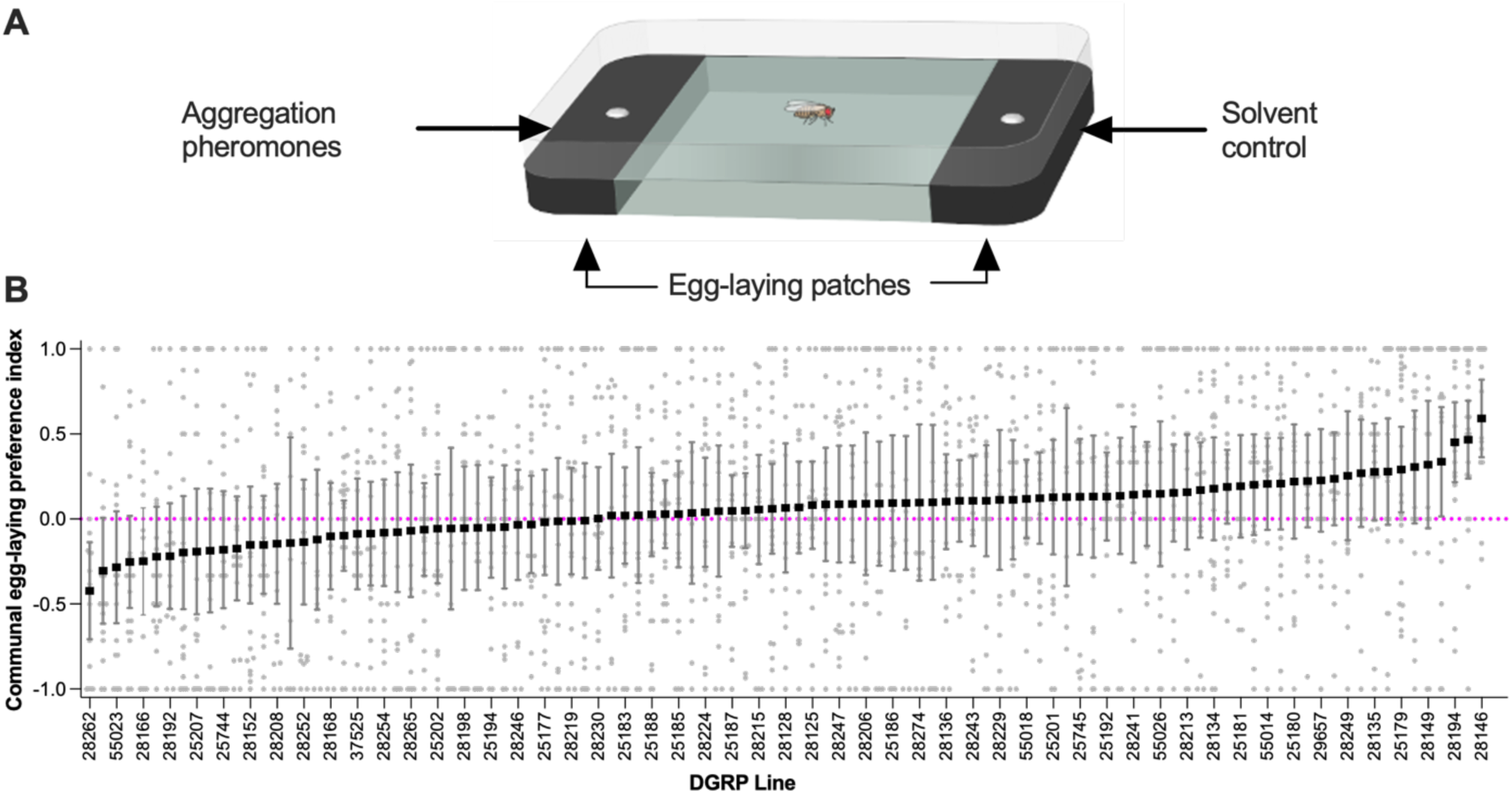
Variation in communal egg-laying preference between DGRP lines. **(A)** Illustration of the communal egg-laying choice assay. A mated female is given an egg-laying choice between two egg-laying patches (black) treated with either aggregation pheromones to mimic communal egg laying or a solvent control. **(B)** Average communal egg-laying preference index in DGRP lines. A positive index indicates preference for communal egg-laying, whereas a negative index indicates isolated egg-laying. Lines are ordered from lowest to highest attraction to lay eggs communally. The magenta dotted line represents no preference between both patches. Replicates ranged from 15 to 29 per line. Error bars indicate 95% confidence interval. For full statistical analysis, see **Table S2**.

### Sociability assay 2: Communal egg-laying latency

Six DGRP virgin females were mated with 6 *OR* males at 25°C and treated as for the Egg-laying site choice assay. For each assay, a single focal mated female was placed in a group with 5 *OR* males in an egg-laying Petri dish (35×14 mm) containing black food patch (23×1 mm) allowing visualization of the white eggs. To force females to lay eggs on the food patch surface and not on the edge, a liquid 3.5% agar solution was poured in the dish up to the level of the food patch surface. Since both males and females trigger an egg-laying advancement in a focal female (Bailly et al., 2023), male group members were used in this assay to ensure that the first laid egg is from the focal female. To avoid bias induced by male group members, we used males from the same strain (*OR*) for test with all DGRP lines. Egg-laying dishes and flies were housed in a stainless-steel enclosure [63(D) × 71(H) × 120(L) cm] for 24 h at 25 °C. To generate 12:12 LD condition, the chamber was lit internally by white LED lights during the light phase of the day and red (> 620 nm) LED lights during the dark phase. Experiments were started at ZT 5 (14:00 our time) during the light phase of a 12:12 LD cycle. To quantify egg-laying, pictures of the egg-laying dishes were taken with cameras (EOS 1300D Canon, equipped with EF-S 18-55mm III lens) at 15-minute intervals during 24 h (using time-lapse software DigiCamControl). Onset of egg-laying of each DGRP female was manually determined from these pictures using the software ImageJ 1.52a (Schneider et al., 2012).

### Sociability assay 3: Spontaneous social interactions

In the spontaneous social interactions assay, four DGRP male flies were placed at room temperature into an aluminum framework (inner dimensions: 66×30×3mm) closed by transparent glass plates placed on top and bottom of the arena and placed on a white LED background lighting. To prevent an effect of day and time of day, trials were randomly spread out over multiple days with a maximum of two replicates per DGRP line per day. After one minute of acclimatization, freely moving flies were recorded for 10 minutes with a Raspberry Pi Camera Module V2 or a Raspberry Pi High Quality Camera with a 16mm 10MP lens, 30 frames per second capture and high resolution, connected to a Raspberry Pi 4 computer. Aluminum and glass surfaces were cleaned with ethanol to avoid contaminations and disrupting odors between experiments.

All behavioral videos were processed using TGrabs and Trex(Walter & Couzin, 2021). TGrabs was used for initial segmentation and detection of individual flies in each frame, while TRex performed multi-object tracking and maintained individual fly identities across frames. Tracking outputs were only retained if data were available for at least 98% of all frames in a 10-minute video, ensuring high-confidence trajectories for all flies.

To prepare the data for downstream analyses, a custom Python script (available at: https://github.com/jcbilleter-hub/flysociabilitytracker) was used to clean and standardize the tracking output. This script first handled missing data points by interpolating x–y positions and angles. For missing x–y coordinates, gaps shorter than five frames (∼0.17 seconds at 30 fps) were filled using nearest-neighbor interpolation (Downs et al. 2012), as displacement during such short intervals is minimal. Gaps between 5-15 frames (∼0.17–0.5 seconds) were filled using cubic spline interpolation(Tremblay et al., 2006), which maintains trajectory smoothness and better approximates natural fly motion. Gaps longer than 15 frames were excluded from analysis to avoid overfitting or spurious path reconstruction. Angular data (fly orientation) were always interpolated using splines to preserve smooth transitions and avoid abrupt heading changes that would bias calculations of angular velocity or body alignment. From the cleaned trajectories, multiple behavioral parameters were computed using additional custom Python scripts (available at: https://github.com/jcbilleter-hub/flysociabilitytracker). Four social interaction metrics were extracted: (1) frequency of social interactions—the number of interaction bouts between any pair of flies; (2) total duration of social interactions—the cumulative time flies spent interacting; (3) interindividual distance—the average distance between all flies across frames; and (4) nearest-neighbor distance—the mean distance between each fly and its closest conspecific per frame. Social interactions were defined based on established criteria (Schneider et al., 2012): two flies were considered to be interacting if (a) the angle between the focal fly’s body axis and the partner’s centroid was less than 90°, (b) the distance between centroids was ≤2 body lengths, and (c) these conditions persisted for at least 1.5 seconds. These metrics were computed across all four flies in a 10-minute video. To assess locomotor activity, average walking speed (cm/s) over the entire trial was calculated for each fly. Additionally, we identified the central region of the arena to facilitate quantification of exploration. The center was arbitrarily defined as the area bounded by trimming ¼ of the arena’s short side from each edge, forming an inner rectangle. Flies were considered to be in the center when their centroid coordinates fell within this region. Explorative behavior was measured as the percentage of time each fly spent in the defined center region of the arena. All derived metrics were then aggregated for further analysis. Prior to testing all DGRP flies, males and virgin females from three DGRP lines were tested in the spontaneous social interaction assay. No significant differences were observed between males and females in the number of interactions (Fig. S3A), total duration of interactions (Fig. S3B), nearest-neighbor distance (Fig. S4A) and interindividual distance (Fig. S4B). Based on these preliminary results, only males of the DGRP lines were used for convenience.

### Statistical analysis

All statistical analyses were performed using R (version 3.6.3, R Core Team 2020), unless otherwise specified. Generalized linear mixed models (GLMMs) were fitted using the glmmTMB package (Brooks, Mollie et al., 2017)to assess the effects of genotype (DGRP line) on behavioral traits, with *Replicate* or *Date of experiment* included as random intercepts where applicable. Model validation was conducted using the DHARMa package(Hartig, 2025), including simulated residual diagnostics and formal tests for overdispersion and zero inflation (Fig. S3A–G).

For spontaneous social interaction traits, distributions were selected based on empirical data distributions and model fit diagnostics. Frequency of interactions (Fig. 3B) were analyzed using a zero-inflated negative binomial GLMM, while duration of interactions, inter-individual distance, and nearest-neighbor distance (Fig. 3C–E) were modelled using Tweedie GLMMs with a log link due to their continuous, right-skewed, and zero-inflated nature. Activity measures including speed (Fig. 5A) and exploration (i.e. percentage of time spent in the center; Fig. 5B) were also modelled using Tweedie GLMMs. Genotype effects were tested by comparing full models (including DGRP line) to null models using likelihood ratio tests (Chi-square).

Egg-laying behavior was analyzed using multiple GLMMs: a Gamma GLMM was used for egg-laying latency (Fig. 2B), and a binomial GLMM for the phase of the first egg (Fig. 2C). The egg-laying site preference index (Fig. 1B) was analyzed using a quasi-binomial GLMM with overdispersion correction, and weighted quasi-binomial models were used when appropriate. Total egg counts in both latency and site preference assays (Figs. 2E, S1A, S2A) were modelled using quasi-Poisson GLMMs. To test for main and interaction effects of genotype and social context on egg-laying latency, Type II Wald chi-square tests and a type III two-way ANOVA were used (Fig. 2E).

**Figure 2:**
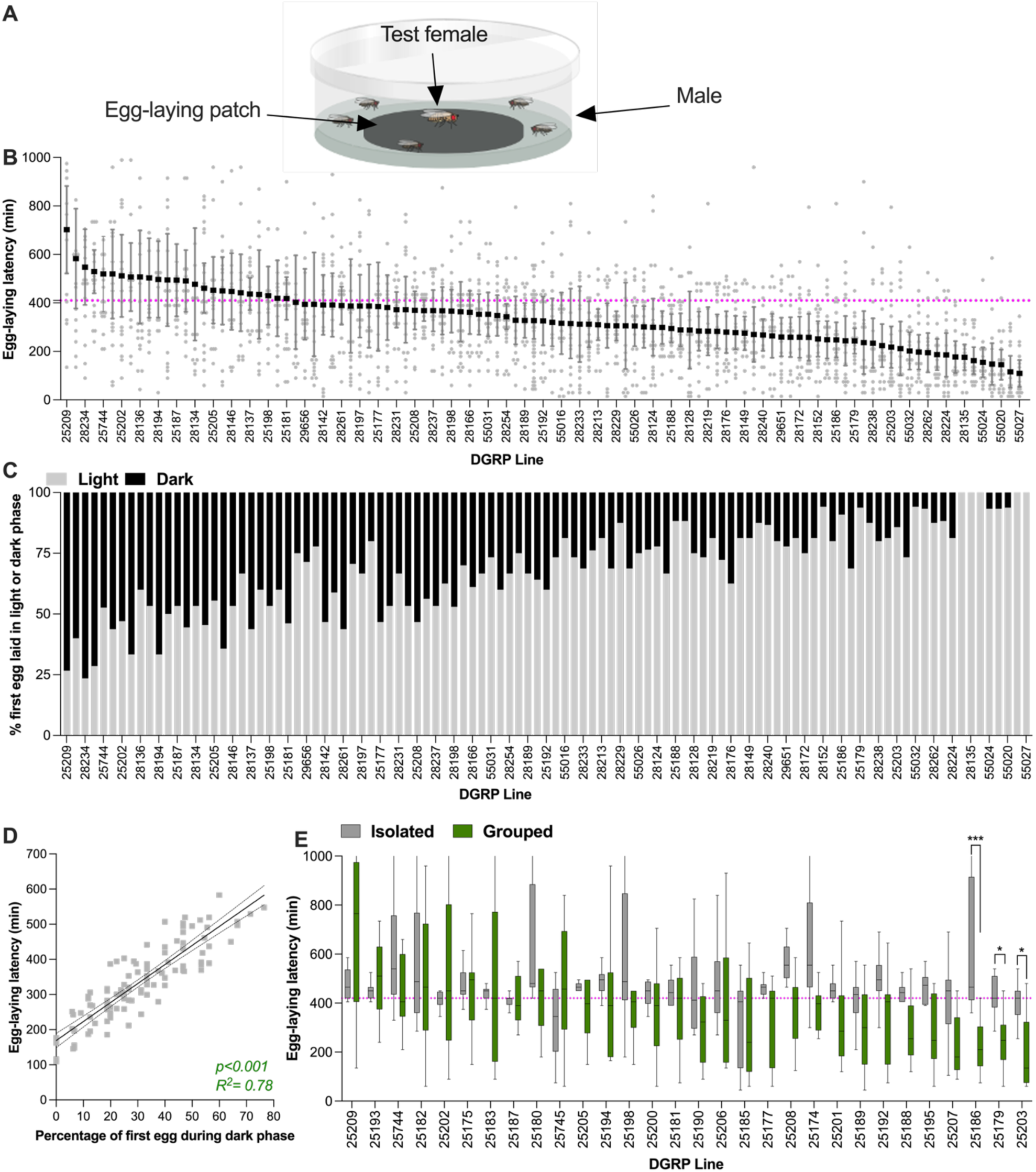
Variation in egg-laying latency in group in DGRP lines. **(A)** Illustration of the egg-laying latency assay. A focal mated DGRP female was placed with 5 *OR* males in a circular dish containing an egg-laying substrate. Egg-laying latency in a group is measured as the time a female takes to lay her first egg after introduction to the dish. **(B)** Average egg-laying latency of DGRP female in group, organized from longer (left) to shorter (right) latency. Because the assay lasts 24hrs and is placed in a 12:12LD cycle, the assay spans light and dark phase. The dotted magenta line represents the onset of the dark phase, below is time in the light phase. **(C)** Percentage of flies per DGRP line that laid their first egg during the light (grey) vs during the dark (black) phase. DGRP lines are ordered as in B. **(D)** Scatterplots and linear regression between mean egg-laying latency per line and percentage of first egg during the night for the 105 DGRP lines. **(E)** Average egg-laying latency by a DGRP mated female isolated (grey, means in black) or grouped, (green, means in black). The dotted magenta lineindicates the onset of darkness. Groups joined by asterisk indicate differences between isolated and grouped as indicated by a post-hoc test (*:<0.05; ***:<0.001). DGRP lines are ordered based on the data in B. Replicates per genotype ranged from 15-25. Error bars indicate 95% confidence interval. For full statistical analysis, see **Table S2.**

All variables used in correlation models were *z*-scored (mean-centered and scaled to unit variance) prior to modelling. For behavioral correlations within and between assays, linear regression analyses were performed in GraphPad Prism 10 (GraphPad Software, La Jolla, CA), and are reported as slope ± SE, *F*-statistic, *p*-value, and *R*² (Figs. S1B, 2D, S2B, 4A–F, 5C–J, 6A–D). For selected comparisons between specific DGRP lines (Fig. S3A–D), Mann–Whitney U tests were performed in GraphPad Prism and followed by false discovery rate (FDR) correction. A full list of statistical tests used, along with response variables, explanatory factors, test statistics, and figure panel references, is provided in Supplementary Table S2.

### Genome Wide Association Study

A genome wide association study (GWAS) was performed for four quantified traits: egg-laying site choice, egg-laying latency, nearest-neighbor distance and duration of social interactions. The genotype data from the DGRP Freeze 2 was obtained in binary PLINK format from the DGRP2 website (http://dgrp2.gnets.ncsu.edu/data.html). Quality control of the genotype data was done using PLINK (v2.0). We filtered for minor allele frequency (>0.01), missing call rate (<0.15) and genotype missingness (<0.1). Linkage disequilibrium pruning was performed with the independent pairwise command, with a window size of 50, a step size of 5 and an r^2^ threshold of 0.5. The filtering resulted in the inclusion of 1,020,854 variants and 105 DGRP lines. A linear mixed model (LMM) was fitted using the GMMAT R-package (H. Chen et al., 2016). *Wolbachia* infection status and the five major inversions *(In(2L)t, In(2R)NS, IN(3R)K, In(3R)P*, and *In(3R)Mo)* were included as fixed-effect covariates. To control for cryptic relatedness, a principal component analysis (PCA) was performed on the variance-standardized genomic relationship matrix, and the top principal components were included as additional covariates. The number of included principal components was based on the genomic inflation factor (λ). Population structure was further accounted for by including the genomic relationship matrix as a random effect in the model. Resulting *p-*values were corrected for multiple testing using the Benjamini-Hochberg procedure to control for false discovery rate (Benjamini & Hochberg, 1995).

## Results

### Variation in communal egg-laying preference

To investigate potential variation in sociability among DGRP lines, we began by quantifying preference for communal versus isolated egg-laying. Communal egg-laying is activated by aggregation pheromones deposited by mated females on egg-laying sites (Duménil et al., 2016; Verschut et al., 2023a). We tested this behavior by providing single mated females with a choice between laying eggs on a food patch treated with mated female pheromones or on a solvent-treated control patch (Fig. 1A). We quantified each DGRP line’s preference for communal egg-laying by calculating a preference index based on the number of eggs laid on the pheromone-treated (communal) versus solvent-treated (isolation) side. A higher preference for the pheromone-treated patch (i.e., a positive egg-laying site preference index) indicates greater sociability, whereas a preference for the solvent-treated patch indicates lower sociability.

Preference for communal egg-laying varied significantly among DGRP lines (Fig. 1B; p<0.001), with an average preference index of 0.052, signifying no preference for laying eggs near aggregation pheromones on average over the lines, and line-specific means ranging from -0.42 to +0.59. These results indicate that sociability in the DGRP population spans a wide spectrum, from actively preferring communal egg-laying to actively avoiding it, revealing opposite sociability tendencies. Since this sociability measure is based on egg production, it could be confounded by differences in fecundity between lines. To assess this, we quantified the number of eggs laid within 24hrs for each DGRP line (Fig. S1A) and examined its correlation with the communal egg-laying preference index (Fig. S1B). Fecundity varied significantly among DGRP lines (Fig. S1A; p<0.001), but this variation did not correlate with preference for communal egg-laying (Fig. S1B; R²=0.001). We conclude that fecundity differences do not confound the interpretation of communal egg-laying preference.

### Variation in communal egg-laying latency

Another aspect of sociability in *Drosophila* is the observation that females from certain wild-type strains lay eggs faster in the presence of other flies compared to when isolated, while other strains lay eggs at the same latency when isolated or in a group (Bailly et al., 2023). We investigated the extent of variation in this behavior amongst DGRP lines by quantifying egg-laying latency of a focal DGRP female grouped with 5 males (Fig. 2A). Egg-laying latency in groups varied significantly among DGRP lines (Fig. 2B; p < 0.001), with an average egg-laying latency of 339 minutes and means per line ranging from 109 to 702 minutes.

Egg-laying advancement in grouped compared to isolated females was originally discovered in females from a wild-type *D. melanogaster* strain called *Oregon-R.* These females wait for dark conditions to begin egg-laying when isolated, but commence laying eggs under light conditions when in a group (Bailly et al., 2023). In this strain, social context takes precedence over light conditions. As laying eggs during the day is a response to being in a group and, thus, a measure of sociability, the percentage of flies that laid eggs during the day was also quantified for each of the 105 DGRP strains. This revealed significant variation in which phase (light or dark) females began egg-laying (Fig. 2C; p<0.001). To investigate whether egg-laying latency and probability to lay eggs in the light or dark phase may quantify a similar trait, the average egg-laying latency in a group per line was correlated with the percentage of females that laid their first egg during the night for a given DGRP. The strong correlation between these two measures (Fig. 2D; R² = 0.78) suggests they quantify a similar process and that laying eggs in groups during the light phase indicates high sociability, whereas laying eggs during the dark phase indicates lower sociability. This hypothesis would be supported if females that laid eggs during the day in groups, would only lay eggs at night when isolated - then egg laying during the day can be considered a direct reaction to being in a group. To test this, we analyzed the egg-laying latency and light phase of isolated vs grouped flies in 30 of the 105 DGRP lines. Isolated females laid eggs primarily at night in 23 of the 30 lines, whereas in groups the females in these strains mostly started laying earlier (Figure 2E). Thus, these lines tended to respond with egg-laying advancement in a social context. The 7 remaining lines did either not alter their egg-laying latency in a group compared to being isolated, or even appear to delay their egg-laying in a group. Of these 7 lines, only 2 lines clearly lay their eggs during the day as isolated females. These observations indicate that laying eggs during the day is mostly a property of grouped flies, but not in all strains. There was a significant overall effect of social context (Fig. 2E; p< 0.001) and differences between lines (Fig. 2E; p<0.0001), confirming our hypothesis that the behavior of grouped flies differ from isolated flies. There was however a strong interaction between social context and genotypes, reflecting the range of responses from egg-laying advancement to delaying in a group (Fig. 2E; p<0.001). Overall, we conclude that most lines that lay eggs during the day in a group respond to their social group and are thus sociable, while lines that lay eggs at night do not respond to their social environment and are thus not sociable in this assay.

This sociability metric, tied to egg production, could be influenced by fecundity variation between lines. To assess fecundity’s impact on egg-laying latency, 24-hour egg counts were tallied for each DGRP line (Fig. S2A), and tested for correlation with egg-laying latency (Figure S2B). Fecundity varied significantly between DGRP lines (Fig. S2A; p < 0.001). However, this inter-line variation did not correlate with egg-laying latency (Fig. S2B; R² = 0.018) indicating that fecundity does not confound the interpretation of egg-laying latency results.

### Variation in spontaneous social interactions

Social distance is a well-established proxy of sociability that translates between species (Anderson et al., 2017; Buijs et al., 2011; Fernandez et al., 2017; Lough et al., 2015; White & Chapman, 1994). Shorter social distance is itself presumably a proxy for a higher number and duration of social contacts, which all indicate stronger attraction to others and thus higher sociability. Metrics such as nearest-neighbor and interindividual distance make use of social distance to respectively quantify the average shortest distance between a given individual and its closest neighbor, and the average spacing between individuals within a group. We used automated behavioral tracking to simultaneously quantify number and duration of social interactions as well as interindividual distance and nearest-neighbor in DGRP lines. Because the four measure are done simultaneously on the same individuals, we also explored correlations between these traits in order to establish whether they represent the same dimension of sociability.

These sociability traits were quantified in a spontaneous social interactions assay, where three male flies from the same DGRP lines were placed into a rectangular arena where they were left for 10 minutes to freely interact (Fig. 3A). Only male DGRP flies were used, as preliminary assays in three genetically distinct lines (including both inbred and outbred strains) revealed no significant sex differences in sociability measures (Fig. S3).

**Figure 3:**
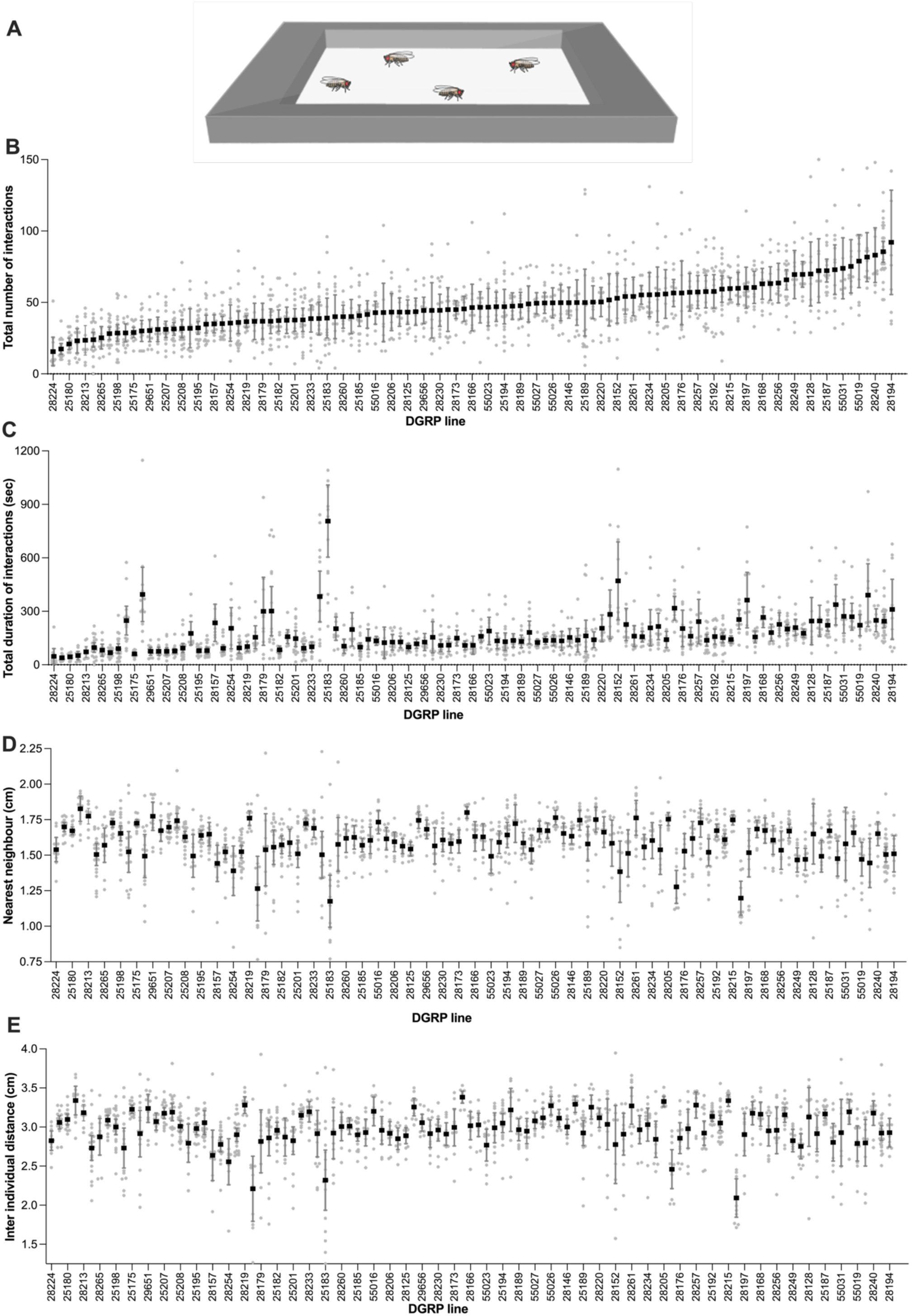
Social distance and interactions vary continuously in DGRP lines. **(A)** Illustration of the spontaneous social interactions assay. Four males from the same DGRP line were placed in a rectangular arena and number of interactions, total duration of all interactions, nearest neighbor distance, inter individual distance, speed and exploration were quantified during 10 minutes. **(B)** Total number and **(C)** duration of all social interactions among 4 flies from the same DGRP line. **(D)** Average nearest-neighbor distance and **(E)** Average interindividual distance between 4 flies from the same DGRP line. DGRP lines are ordered on low to high sociability based on the data in **(B)**. For each DGRP line, between 9 and 27 replicate assays were conducted (overview of replicates per DGRP line in Table S1). Error bars indicate 95% confidence interval. For full statistical analysis and methods, see **Table S2**.

We first quantified social interactions in the DGRP lines. The number of interactions varied significantly among DGRP lines (Fig. 3B; p<0.001), with an average total number of interactions of 47 per 10 minutes in the DGRP populations, ranging from 16 to 92 per line. The total duration of these interactions also varied significantly (Fig. 3C; p<0.001) as well as nearest-neighbors (Fig. 3D; p<0.001) and inter-individual distance (Fig. 3E; p<0.001).

We next determined correlation between these four traits quantified on the same individuals to explore if they may capture a similar dimension of sociability. Total number of social interactions and their total duration are correlated (Fig. 4A), suggesting that these two measures based on social interactions between two individuals represent a similar dimension of sociability. Interindividual distance and nearest neighbor are also strongly correlated (Fig. 4B), suggesting that measures based on distance between individuals represent a similar dimension of sociability. This indicates that the probability of a fly being close to another is the highest when the whole group is more closely clustered. However, sociability measures based on social distance (interindividual distance & nearest-neighbor) are not correlated with those based on number of social interactions (Fig. 4C-D), but are anti-correlated with those based on duration of interactions (Fig. 4E-F). Taken together these results show that sociability varies in a least two different dimensions within one sociability paradigm: social distance and social interactions.

**Figure 4:**
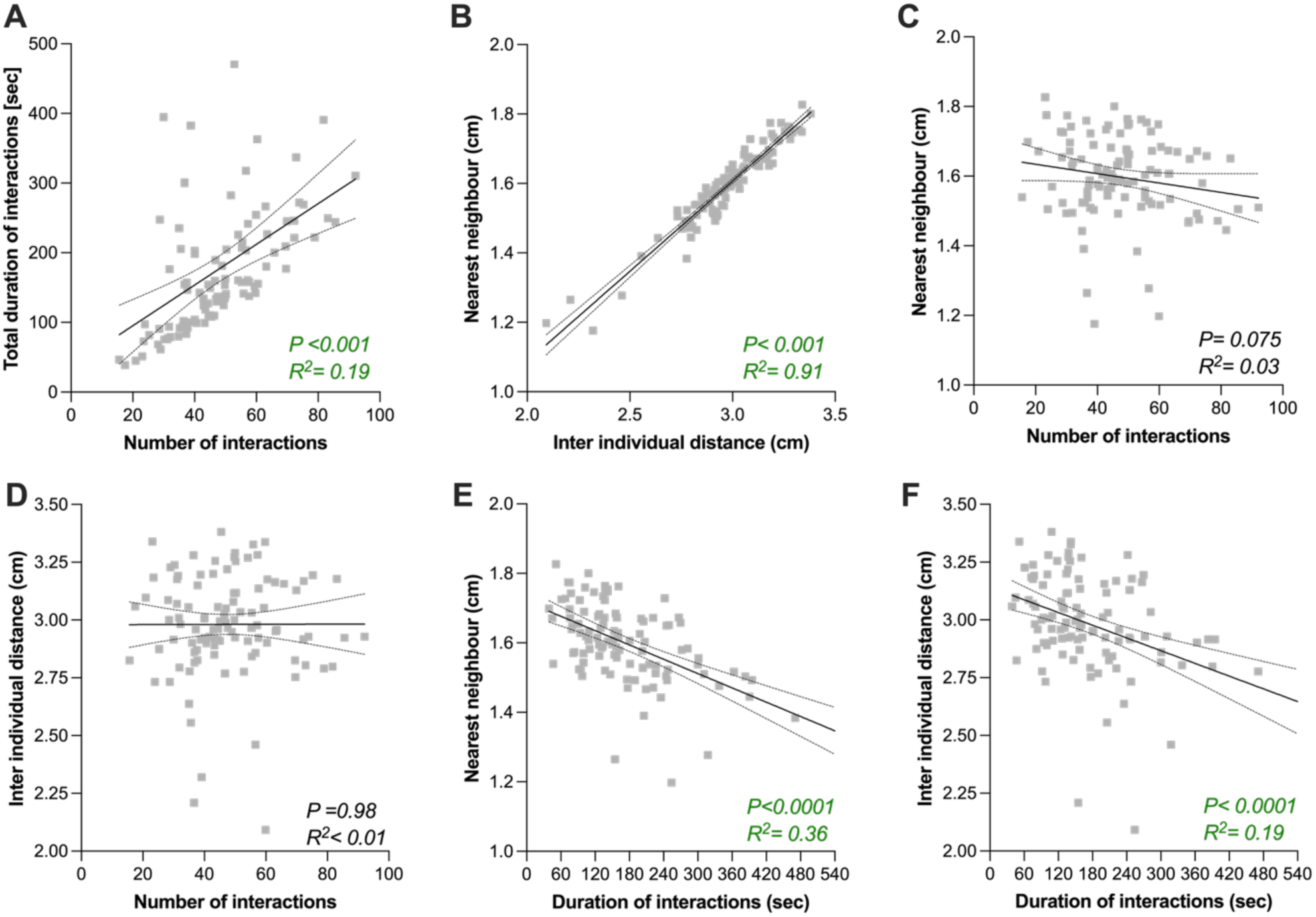
Correlation between sociability metrics in the spontaneous social interactions assay. (A-F) Scatterplots and linear regression between total number of interactions and total duration of all interactions for the 105 DGRP lines. The continuous black line indicates the slope and dashed lines 95% confidence interval. Linear regressions between two indicated traits were performed on the mean value for each line. Probability of a significant correlation (P) and the % correlation between the two indicated traits (R^2^) are indicated in each panel. For full statistical analysis, see **Table S2.**

Quantification of social distance and social interactions may be confounded by between-line differences in locomotor activity and reluctance to entering the center of the arena (centrophobism), reducing the area of the arena in which flies may interact. These traits were determined using automated behavioral tracking and correlated with sociability measures to assess their influence (Fig. 5C-J). Average speed varied significantly among DGRP lines (Fig. 5A; p<0.001) as well as time spent in the center of the arena (Figure 5B; p<0.001). Speed did not correlate with the number of social interactions (Fig. 5C), but did correlate with duration of social interactions, and both inter individual and nearest neighbor distances (Figure 5D-F). Time spent in the centre of the arena modestly correlated with social interaction number and duration (Figs. 5G&H), but had a strong correlation with social spacing (Figs. 5I&J). Together, these correlations indicate that measures based on social interactions are modestly influenced by interline differences in speed and centrophobism, but that measure based on social spacing are strongly confounded by speed and centrophobism.

**Figure 5:**
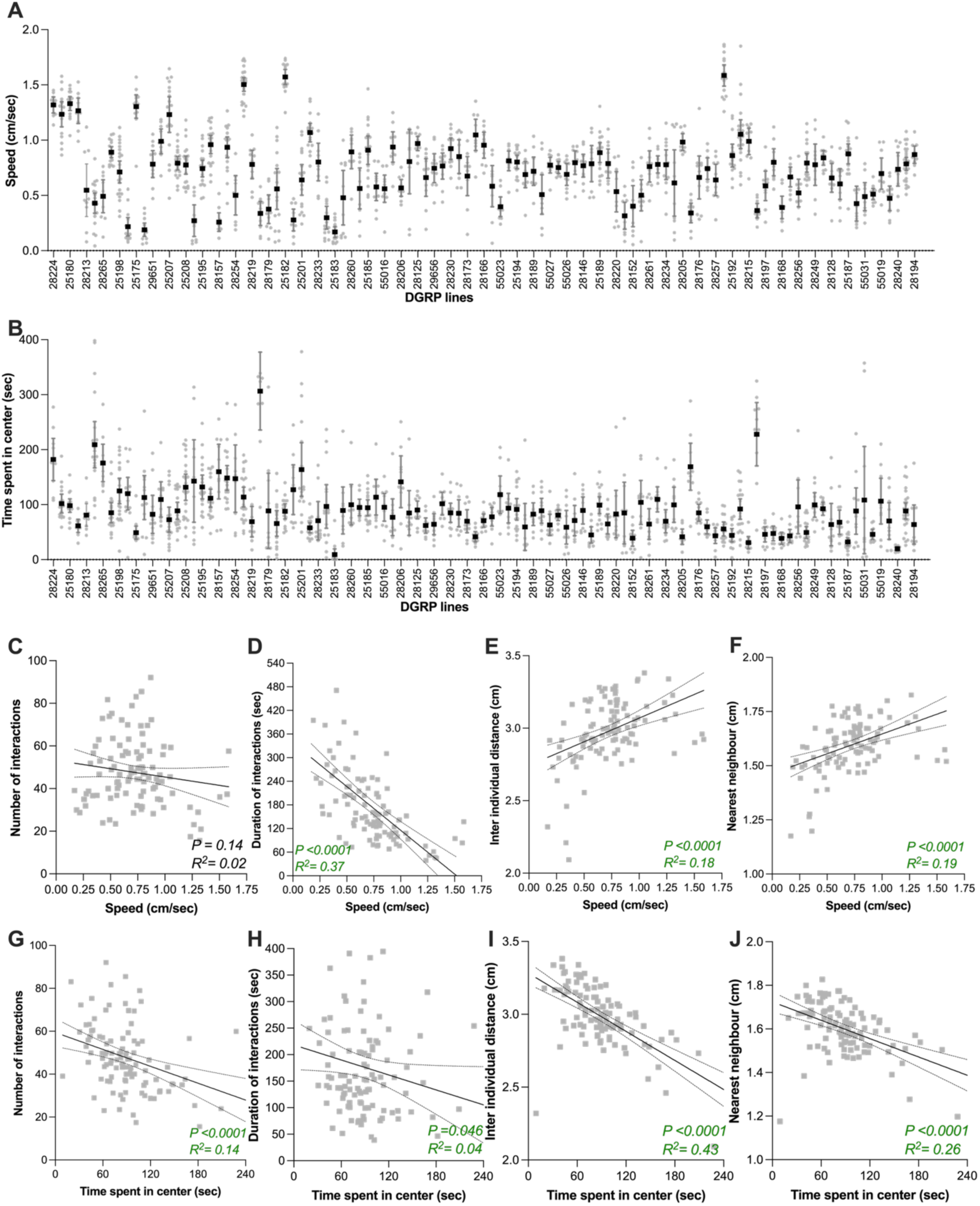
Correlation between sociability metrics and both speed and centrophobism. **(A)** Average speed and **(B)** time spent in the center of the arena of 4 flies from the same DGRP line. DGRP lines are ordered on low to high sociability based on the data in (3B). (**C-J)** Scatterplots and linear regression between total number of interactions and total duration of all interactions for the 105 DGRP lines. Black line indicates the slope and dashed lines 95% confidence interval. Linear regressions between two indicated traits were performed on the mean value for each line. Probability of a significant correlation (P) and the correlation between the two indicated traits (R^2^) are indicated in each panel. For full statistical analysis, see **Table S2.**

### Sociability measures are not correlated

Our data revealed significant phenotypic variation among the 105 DGRP lines tested in traits associated with sociability. We next tested if correlations existed between quantitative performance of a line in one assay and another, in order to test whether variation in sociability translates between different assays and thus represent similar dimensions of sociability. No significant correlations were found between average values in the communal egg-laying latency assay and in the communal egg-laying site preference assay (Fig. 6A), nor between average values in the communal egg-laying preference assay and in the spontaneous social interactions assay (Fig. 6B) nor between average values in the communal egg-laying latency assay and the number of social interactions in the spontaneous social interactions assay (Fig. 6C). This indicates that across all the DGRP lines tested, there was no consistent pattern of lines that scored high for sociability in one of the assays and also scored high for sociability in the other assays.

**Figure 6:**
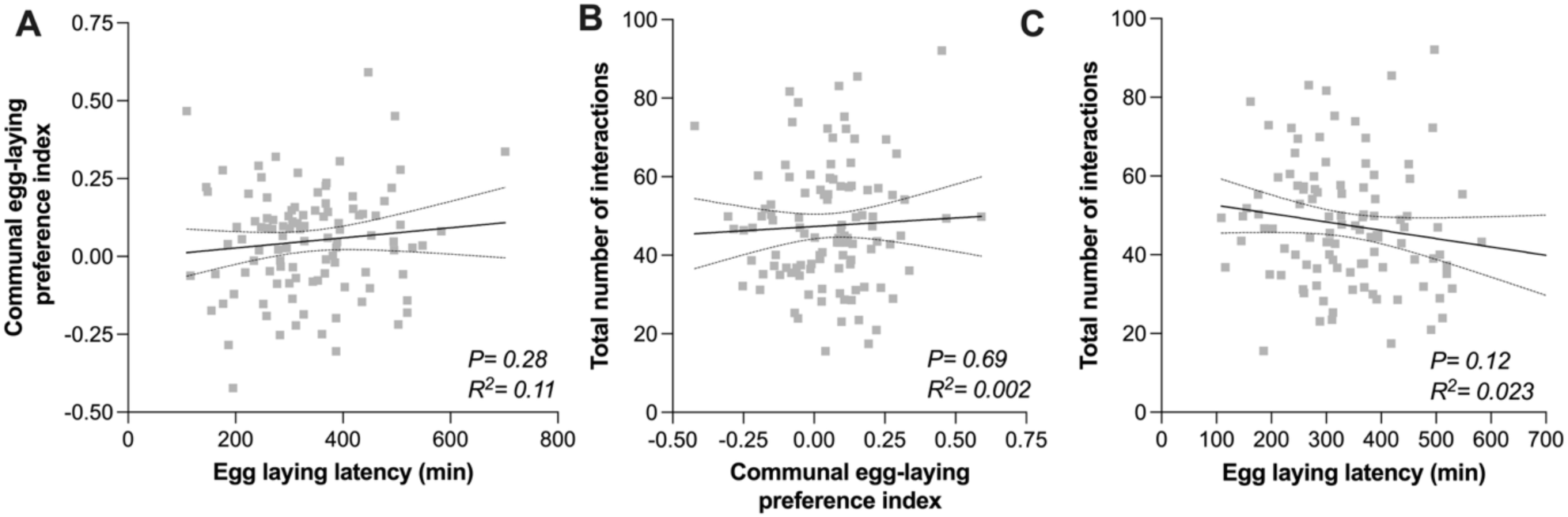
Correlations between the sociability measures of the three behavioral assays. **(A)** Scatterplots and linear regressions between the egg-laying latency and the communal egg-laying preference index for each DGRP line. **(B)** Scatterplots and linear regression between the communal egg-laying preference index and total number of interactions for each DGRP line. **(C)** Scatterplots and linear regression between the egg-laying latency and total number of interactions for each DGRP line. Error bars indicate 95% confidence interval. For full statistical analysis and methods, see **Table S2.**

### Genome Wide association Studies on sociability measures

Significant differences in each of sociability paradigm between DGRP lines suggest genetic variance for the different aspects of sociability. Given that the genome of all DGRP lines have been sequenced, we performed a GWAS on the mean trait values for egg-laying latency (Fig. 2), egg-laying site choice (Fig. 1), duration of social interaction (Fig. 3C) and nearest-neighbor (Fig 3D) to determine how the observed phenotypes can be explained by the genotype. No SNPs, deletions, or insertions reached the genome-wide significance threshold of *p*<10^-6^ in the DGRP GWAS. A list of potentially interesting SNPS (*p* value <10^-1^) are shown in Table S3. A key message here is that there is no overlap between the SNPs associated with the 4 independent GWASs of the different traits.

## Discussion

In this study, we explored sociability in *D. melanogaster* using a multidimensional approach that combines three sociability paradigms, each targeting distinct aspects of an individual’s propensity to interact and respond to others. By testing 105 DGRP lines, we uncovered significant variation among genetically distinct wild-type strains in their responses to conspecifics, thereby demonstrating that sociability in fruit flies varies between individuals or genotypes and exists along a spectrum. Importantly, sociability measures and ranking of the tested lines did not correlate across the three independent paradigms, indicating that sociability cannot be captured in a single trait and instead reflects a set of independent behavioral dimensions.

### Sociability as a multi-dimensional trait

The multidimensional nature of sociability in *D. melanogaster* sheds light on its underlying mechanisms. It likely reflects the diversity of sensory inputs required to mediate social interactions. Spontaneous social interactions involve tactile and chemical cues exchanged through leg-to-leg contact, as well as spacing behaviors modulated by vision, olfaction, gustation, and hearing (Burg, Langan, and Nash, 2013; Jiang et al., 2020; Ramdya et al., 2015; Schneider et al., 2012; Simon et al., 2012) Communal egg-laying relies primarily on olfactory detection of pheromones (Billeter & Wolfner, 2018; Duménil et al., 2016; Verschut et al., 2023a). By contrast, egg-laying latency in the presence of groups depends on visual processing via motion detection (Bailly et al., 2023). Sociability in fruit flies is expressed through multiple behavioral dimensions underpinned by distinct sensory mechanisms, which may explain why variation in one context does not predict variation in another. Our results therefore emphasize the need to consider specific sensory modalities and their contribution to social behaviors when interpreting sociability.

Our results also argue against the existence of a brain centre dedicated to social interactions in *D. melanogaster*. Across multiple assays measuring individual response to others, we observed little consistency in the relative sociability of genotypes. If a central brain centre integrated social cues to guide all social responses, one would expect consistent genotype rankings across assays purported to measure sociability—yet this is not observed. This lack of concordance across contexts suggests that each social paradigm engages distinct biological pathways and neuronal circuits, each with its own genetic and cellular substrates. The concept that sociability cannot be linked to a unique pathway in *Drosophila* finds parallels in the outcome of learning & memory studies in that species. Indeed, testing fruit flies across different memory paradigms reveals no significant correlations in performance across tasks, nor any shared genetic loci (Hamlin et al., 2025). This is not surprising given that different types of memory are viewed as distinct processes that likely depend on different neural circuits (Davis, 2023). From an evolutionary standpoint, such independence would allow each dimension of social behavior to respond to selection pressures with minimal constraints from other traits.

These findings raise a conceptual question: what does “sociability” mean in *D. melanogaster*? In this species, sociability appears multidimensional—expressed through distinct behaviors shaped by different sensory modalities. At the ultimate, evolutionary level, the pronounced phenotypic variation across social paradigms, its heritable component, and its likely effects on fitness imply that each dimension of sociability has been independently shaped by natural and social selection. Consequently, the adaptive value of sociability would have to be examined for each social behavior in its specific ecological and social context as single paradigms may not be reliable proxies for an individual’s overall social tendency. At the proximate, mechanistic level, the apparent independence of social behaviors suggests that distinct neuronal and molecular substrates are engaged in different contexts. Explaining inter-individual variation in sociability therefore would require examining the biological basis of specific social behaviors rather than searching for a unitary sociability pathway or centre.

An alternative framework is to view sociability as a latent state—a hidden variable expressed through distinct observable behaviors depending on context. Under this view, studying a specific social scenario can reveal how social experience modulates particular neuronal circuits or molecular pathways, and how these modulations differ between individuals. The general regulatory principles emerging from such focused analyses, rather than the identity of specific genes or cells, may then apply to other social contexts. By dissecting individual sociability paradigms with genetic, cellular, and physiological approaches, we can identify the mechanisms that define each behavioral dimension and uncover shared principles governing social behaviors. This framework may ultimately clarify both the evolutionary origins and the translational significance of variation in sociability.

### Confounding traits and correlations with activity and exploration

Several caveats must be considered against our data identifying sociability as a multi-dimensional trait. First, the use of inbred lines may reveal phenotypes that are artefacts of inbreeding rather than representative of wild populations. Indeed, a large number of DGRP lines were too sickly to yield reliable sociability measures in our hands. A second explanation for the lack of correlation across assays is that, although sociability may be unidimensional, measurements are influenced by non-social traits unique to each paradigm. Locomotor activity in particular is variable across DGRP lines and closely related to spontaneous social interaction measures. We observed positive correlations between total distance moved and multiple sociability parameters, indicating that activity levels can mask or mimic social behaviors. Previous studies have reported inconsistent associations between activity and sociability in flies (Anderson et al., 2016; Scott et al., 2018), though correlations between locomotion and social behaviors have been observed in other species (e.g., male water striders, *Aquarius remiges*;(Sih et al., 2014)). By contrast, we did not find evidence that low locomotor activity confounded performance in egg-laying assays: lines with the lowest locomotor activity laid eggs on both patches of the arena and showed no reduction in fecundity. Nonetheless, our study highlights the complexity of assigning clear social functions to behaviors when non-social traits such as locomotion or exploration may be involved. Finally, we used females to quantify sociability in both the communal egg-laying and egg-laying latency assays for obvious reasons, but chose to assay males in the social distance assay for logistics reasons. Although our pilot data indicated no significant sex-specific differences, it is very likely that sex by genotype interactions exist in at least some of the DGRP lines. These would have affected correlations between female-specific assays based on egg-laying and non-sex specific ones based on social distance. Nonetheless, the absence of correlation between two closely related female sociability measures—communal egg-laying and egg-laying latency—in DGRP lines reinforces our conclusion that sociability is multidimensional.

### Relevance for understanding sociability in other species

Group formation is an ancient and widespread phenomenon in the animal kingdom. Fossil evidence of aggregation dates back approximately 480 million years (Vannier et al., 2019), suggesting that the propensity to associate with conspecifics may have originated in a common ancestor of extant animals. Consequently, mechanisms promoting social attraction may be conserved to some extent across the animal phylogeny. Identifying a sociability spectrum in *Drosophila melanogaster* adds to growing evidence that variation in social behavior is a basic feature of animal populations, regardless of social complexity. This makes *D. melanogaster* a valuable model for studying the evolution and mechanisms of sociability. However, comparing social behaviors across species remains difficult because behaviors can differ in form while serving similar functions.

The *Drosophila* Genetic Reference Panel (DGRP), which combines extensive natural variation with powerful genetic tools, provides a strong framework for uncovering the genetic basis of social behaviors. Each DGRP line represents a unique genotype from a single natural population, allowing behavioral variation to be linked to genetic polymorphisms using genome-wide association studies. Unfortunately, more than 50 lines were excluded from our analysis due to exceptionally low fecundity and locomotor activity in our hands, substantially reducing statistical power and likely explaining why no SNPs reached genome-wide significance. This represents a missed opportunity, as identifying genes associated with sociability in flies would allow direct comparisons with human sociability loci (Bralten et al., 2021). Given that approximately 75% of human disease-related genes have fly orthologs (Millburn et al., 2016), overlapping genetic associations between flies and humans would suggest that certain molecular influences on sociability are evolutionarily conserved. Such candidate genes could subsequently be validated functionally in flies and in vertebrate models, such as mice, providing a powerful comparative framework across taxa. We have previously demonstrated such an approach by examining a knockout of the dopamine type 2 receptor, observing parallel effects in mice and flies on both the number of social interactions and individual spacing (Ike et al., 2023).

Recognizing that sociability is multidimensional raises the question of whether a composite sociability index should be generated for each DGRP line by integrating scores across behavioral paradigms. This approach parallels strategies used in human studies, where multiple questionnaire-based dimensions of sociability are combined into a single composite score to facilitate genetic analysis (Bralten et al., 2021). Similarly, multidimensional behavioral assays have been applied in *D. melanogaster* to model autism spectrum disorder–related phenotypes (Hope et al., 2019). A multidimensional framework may thus have better captured the complexity of sociability. However, in our study, the absence of correlation between behavioral paradigms indicated that these assays measure distinct, non-overlapping aspects of social behavior. This justified our decision not to collapse them into a single sociability score.

### Conclusions

Although *Drosophila melanogaster* has traditionally been viewed as a solitary species, our results reveal substantial natural variation in how wild-type strains engage with conspecifics. This variability demonstrates that sociability is not limited to overtly social species but can differ widely within animals possessing relatively simple social structures. Such variation may provide the evolutionary substrate for the emergence of more complex social systems.

We further show that sociability in *D. melanogaster* is variable, multidimensional, and genetically influenced. The lack of correlation across behavioral paradigms argues against treating sociability as a single, unified trait and instead supports its interpretation as a spectrum of distinct behaviors regulated by partly independent mechanisms. Future research should aim to integrate genetic, neural, and behavioral analyses to elucidate the mechanisms underlying each dimension of sociability. Comparative studies across taxa will also be essential for determining the extent to which sociability-related genes and pathways are evolutionarily conserved. This multidimensional perspective opens new avenues for the genetic and neural dissection of sociability in flies and strengthens their value as a comparative model for understanding how social behavior evolves and is regulated across animals.

## Acknowledgement

We thank A. Bailly, J. Wagenaar, and R. Rajadhyaksha for help collecting phenotypic data on DGRP lines. We are grateful to Apu Ramesh for critical discussions on early versions of the manuscript. This work was supported by Adaptive Life scholarship from the Faculty of Science and Engineering (FSE) of the University of Groningen to TPMB. and SJCL. We are also grateful to the FSE for funding a short post-doctoral stay for AS.

## Author contributions

TPMB, SL, RE, BW and JCB designed the experiments. TPMB, SL, MvD and AJ collected the data. MvD developed the communal egg-laying choice assay used in this study. AS and KF developed the behavioral tracking analysis pipeline. AS performed statistical analysis on all behavioral traits. A. performed the GWAS. TB, SL and JCB wrote the manuscript.

## Supplementary Figures

**Figure S1:**
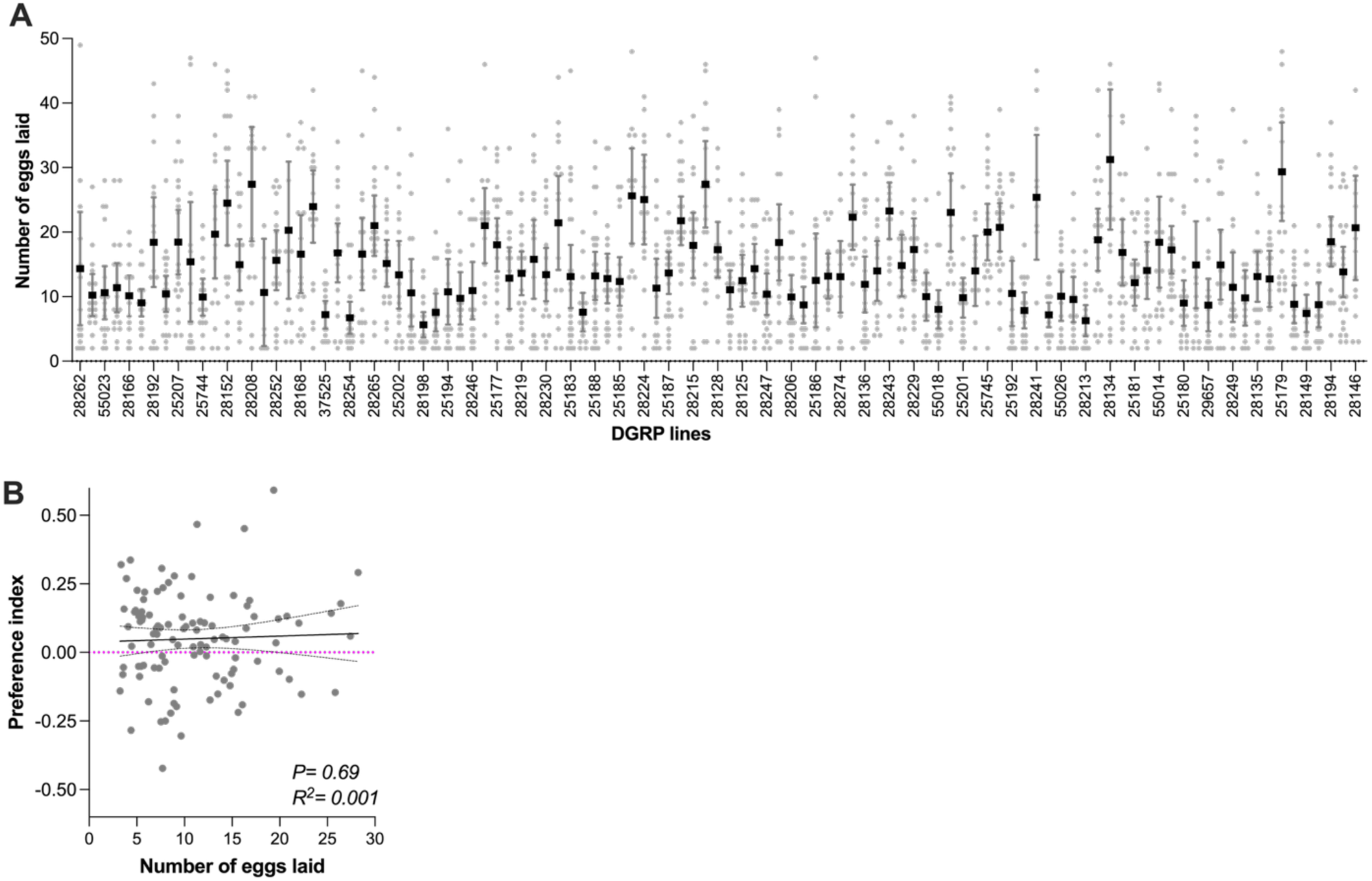
No correlation between communal egg-laying preference and fecundity between DGRP lines. **(A)** Average number of eggs laid in the communal egg-laying preference assay in DGRP lines ordered as in Figure 1B. Replicates ranged from 15 to 29 per line. Error bars indicate 95% confidence interval. (**B**) No correlation between average communal egg-laying preference and number of egg-laid in DGRP lines. For full statistical analysis and methods, see **Supplementary Table S2.**

**Figure S2:**
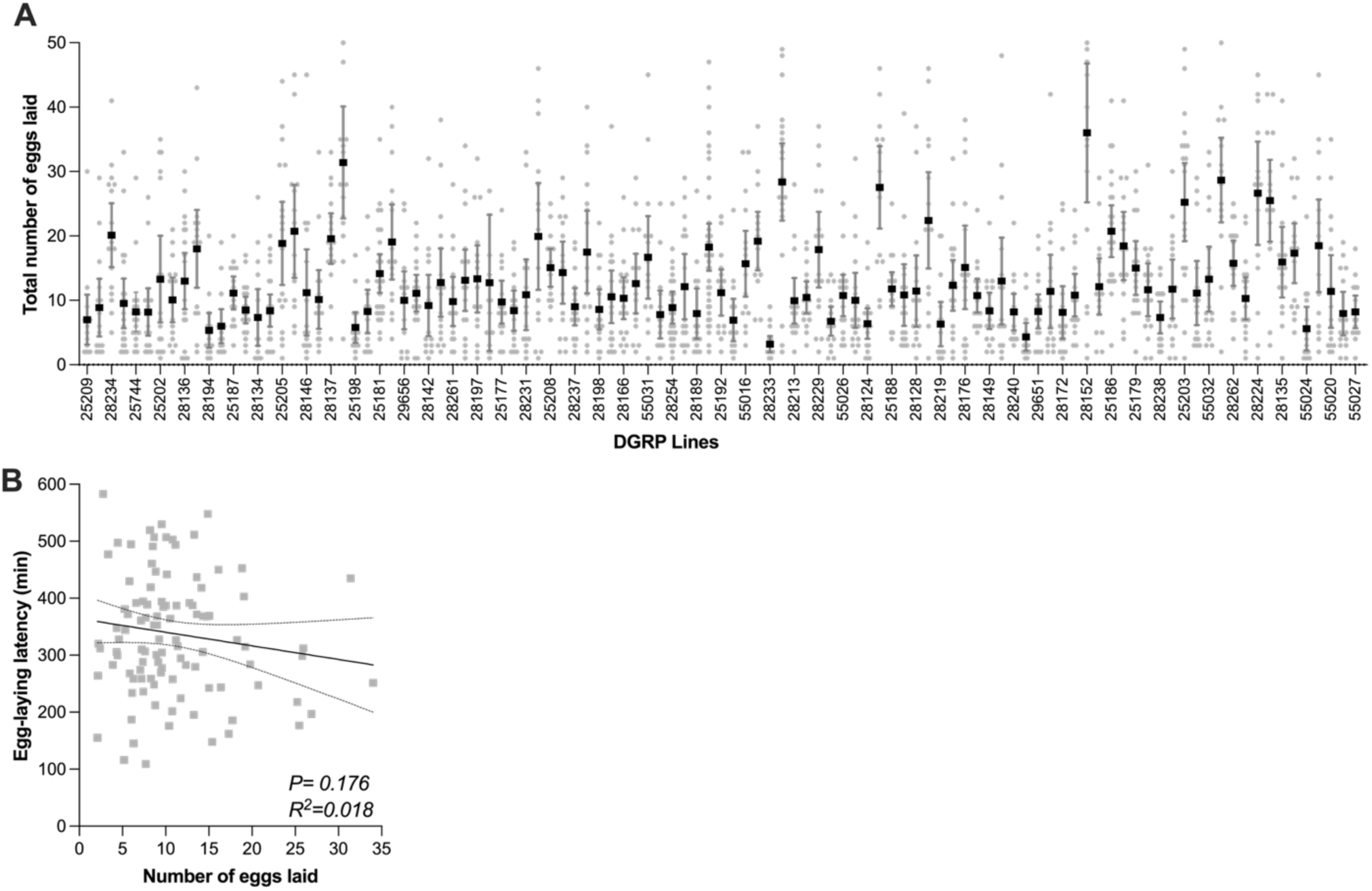
Lack of correlation between egg-laying latency in group and fecundity between DGRP lines. **(A)** Average number of eggs laid in the communal egg-laying assay in DGRP lines ordered as in Figure 2B. Replicates ranged from 15 to 29 per line. Error bars indicate 95% confidence interval. (**B**) No correlation between average egg-laying latency in group and number of egg-laid in DGRP lines. For full statistical analysis, see **Supplementary Table S2.**

**Figure S3:**
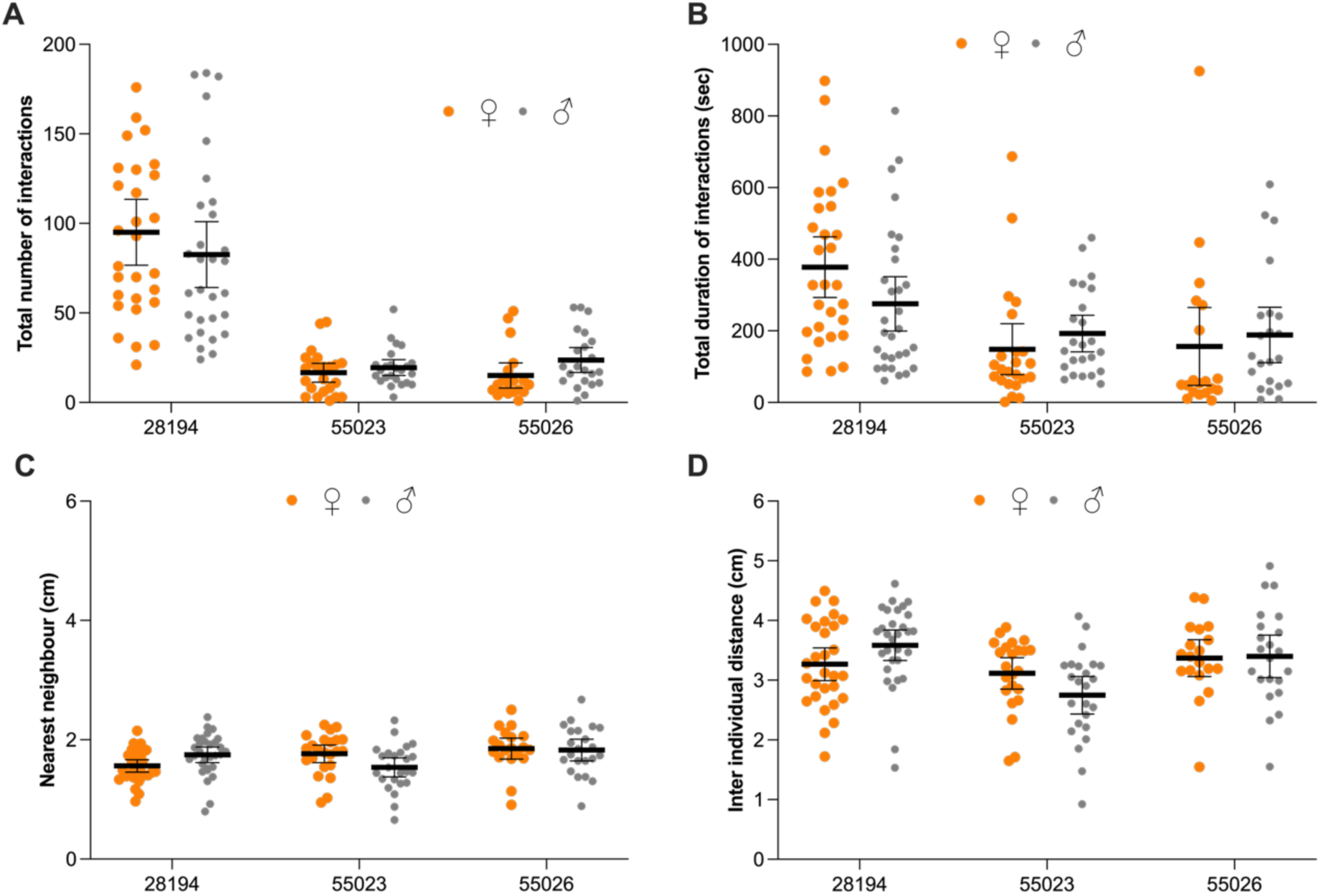
No significant sexual dimorphism in spontaneous social interaction metrics in three DGRP strains. **(A)**Total number, **(B)** duration of all social interactions, **(C)** Average nearest neighbour distance and **(D)** Average inter individual distance between 4 female (orange dots) or male (grey dots) flies from the same DGRP line. For each of the three DGRP lines, between 9 and 27 replicate assays were conducted. Error bars indicate 95% confidence interval. No statistically significant differences were found between males and females in any of the metrics. For full statistical analysis and methods, see **Table S2**.

**Table S1:** Drosophila Genetic reference panels DGRP lines used in this study.

| **DGRP lines** |  |  |
| --- | --- | --- |
| 25174 | 28179 | 55023 |
| 25175 | 28188 | 55024 |
| 25177 | 28189 | 55026 |
| 25179 | 28192 | 55027 |
| 25180 | 28194 | 55031 |
| 25181 | 28197 | 55032 |
| 25182 | 28198 | 83728 |
| 25183 | 28205 |  |
| 25185 | 28206 |  |
| 25186 | 28207 |  |
| 25187 | 28208 |  |
| 25188 | 28213 |  |
| 25189 | 28215 |  |
| 25190 | 28219 |  |
| 25192 | 28220 |  |
| 25193 | 28224 |  |
| 25194 | 28229 |  |
| 25195 | 28230 |  |
| 25198 | 28231 |  |
| 25200 | 28233 |  |
| 25201 | 28234 |  |
| 25202 | 28235 |  |
| 25203 | 28237 |  |
| 25205 | 28238 |  |
| 25206 | 28240 |  |
| 25207 | 28241 |  |
| 25208 | 28243 |  |
| 25209 | 28246 |  |
| 25744 | 28247 |  |
| 25745 | 28249 |  |
| 28124 | 28252 |  |
| 28125 | 28254 |  |
| 28128 | 28256 |  |
| 28132 | 28257 |  |
| 28134 | 28260 |  |
| 28135 | 28261 |  |
| 28136 | 28262 |  |
| 28137 | 28265 |  |
| 28140 | 28274 |  |
| 28142 | 28276 |  |
| 28146 | 29651 |  |
| 28149 | 29656 |  |
| 28152 | 29657 |  |
| 28157 | 37525 |  |
| 28166 | 55014 |  |
| 28168 | 55016 |  |
| 28172 | 55018 |  |
| 28173 | 55019 |  |
| 28176 | 55020 |  |

**Supplementary Table S2:** Summary of statistical analyses organized per figure panels.

| **Fig** | **Test** | **Response variable** | **Explanatory factor** | **Results** | **Post-hoc tests** |
| --- | --- | --- | --- | --- | --- |
| 1B | Quasi-binomial GLMM | Communal egg-laying preference index | DGRP lines | $\chi^2 = 2114.7$, df = 104, $p < 0.001$ | NA |
| S1A | Quasi-Poisson GLMM | Total number of eggs | DGRP lines | $\chi^2 = 513.65$, df = 104, $p < 0.001$ | NA |
| S1B | Linear regression | Communal egg-laying preference index | Number of eggs laid | slope = $0.001 \pm 0.003$ SE, $F(1, 103) = 0.155$, $p = 0.694$, $R^2 = 0.001$ | NA |
| 2B | Gamma GLMM | Egg-laying latency (mins) | DGRP lines | $\chi^2 = 352.41$, df = 104, $p < 0.001$ | NA |
| 2C | Binomial GLMM | % first egg laid in light or dark phase | DGRP lines | $\chi^2 = 237.56$, df = 104, $p < 0.001$ | NA |
| 2D | Linear regression | Egg-laying latency (mins) | Percentage of first egg during dark phase | slope = $5.41 \pm 0.28$ SE, $F(1,103) = 367.5$, $p < 0.0001$, $R^2 = 0.78$ | NA |
| 2E | Type II Wald chisquare test | Egg-laying latency (mins) | Group [isolated, grouped] | $\chi^2 = 25.19$, df = 1, $p < 0.001$ | NA |
| | Type II Wald chisquare test | Egg-laying latency (mins) | DGRP line | $\chi^2 = 97.21$, df = 29, $p < 0.001$ | NA |
| | Type 3 two-way Anova | Egg-laying latency (mins) | Genotype [DGRP line] x Social context [isolated, grouped] | $F(29,10) = 3.10$, $p < 0.001$ | 25179: $p = 0.017$, 25186: $p < 0.001$, 25203: $p = 0.016$ |
| S2A | Quasi-Poisson GLMM | Total number of eggs | DGRP lines | $\chi^2 = 614.71$, df = 104, $p < 0.001$ | NA |
| S2B | Linear regression | Egg-laying latency (mins) | Number of eggs laid | slope = $-2.38 \pm 1.75$ SE, $F(1,103) = 1.86$, $p = 0.176$, $R^2 = 0.018$ | NA |
| 3B | Negative binomial GLMM | Frequency of social interactions | DGRP lines | $\chi^2 = 781.35$, df = 104, $p < 0.001$ | NA |
| 3C | Tweedie GLMM | Duration of social interactions (sec) | DGRP lines | $\chi^2 = 1098.1$, df = 104, $p < 0.001$ | NA |
| 3D | Tweedie GLMM | Inter-individual distance (cm) | DGRP lines | $\chi^2 = 549.71$, df = 104, $p < 0.001$ | NA |
| 3E | Tweedie GLMM | Nearest-neighbour distance (cm) | DGRP lines | $\chi^2 = 575.58$, df = 104, $p < 0.001$ | NA |
| S3A | Mann-Whitney with FDR | Total number of interactions | DGRP lines | 28194: 0.41<br>55023: 0.41<br>55026: 0.07 | NA |
| S3B | Mann-Whitney with FDR | Total duration of interactions (sec) | DGRP lines | 28194: 0.069<br>55023: 0.069<br>55026: 0.219 | NA |
| S3C | Mann-Whitney with FDR | Nearest neighbour distance (cm) | DGRP lines | 28194: 0.026<br>55023: 0.026<br>55026: 0.638 | NA |
| S3D | Mann-Whitney with FDR | Inter-individual distance (cm) | DGRP lines | 28194: 0.113<br>55023: 0.113<br>55026: 0.937 | NA |
| 4A | Linear regression | Total duration of interactions | Number of interactions | slope = $2.92 \pm 0.6$ SE, $F(1,103) = 23.6$, $p < 0.0001$, $R^2 = 0.19$ | NA |
| 4B | Linear regression | Nearest neighbour distance (cm) | Inter individual distance (cm) | slope = $0.52 \pm 0.015$ SE, $F(1,103) = 1068$, $p < 0.0001$, $R^2 = 0.91$ | NA |
| 4C | Linear regression | Nearest neighbour distance (cm) | Number of interactions | slope = $-0.001 \pm 0.00075$ SE, $F(1,103) = 3.23$, $p = 0.075$, $R^2 = 0.03$ | NA |
| 4D | Linear regression | Inter-individual distance (cm) | Number of interactions | slope = $3.306e-005 \pm 0.0014$ SE, $F(1,103) = 0.0006$, $p = 0.98$, $R^2 < 0.01$ | NA |
| 4E | Linear regression | Nearest neighbour distance (cm) | Duration of interactions (sec) | slope = $-0.0006865 \pm 8.974e-005$ SE, $F(1,103) = 58.52$, $p < 0.0001$, $R^2 = 0.36$ | NA |
| 4F | Linear regression | Inter individual distance (cm) | Duration of interactions (sec) | slope = $-0.0009172 \pm 0.0001$ SE, $F(1,103) = 24.4$, $p < 0.0001$, $R^2 = 0.19$ | NA |
| 5A | Tweedie GLMM | Speed (cm/sec) | DGRP lines | $\chi^2 = 1535.1$, $df = 104$, $p < 0.001$ | NA |
| 5B | Tweedie GLMM | Time spent in centre | DGRP lines | $\chi^2 = 901.49$, $df = 104$, $p < 0.001$ | NA |
| 5C | Linear regression | Number of interactions | Speed (cm/sec) | slope = $-7.803 \pm 5.246$ SE, $F(1,103) = 2.212$, $p = 0.14$, $R^2 = 0.02$ | NA |
| 5D | Linear regression | Duration of interactions (sec) | Speed (cm/sec) | slope = $-222.3 \pm 28.45$ SE, $F(1,103) = 61.03$, $p < 0.0001$, $R^2 = 0.37$ | NA |
| 5E | Linear regression | Inter-individual distance (cm) | Speed (cm/sec) | slope = $0.3266 \pm 0.06$ SE, $F(1,103) = 23.04$, $p < 0.0001$, $R^2 = 0.18$ | NA |
| 5F | Linear regression | Nearest neighbour distance (cm) | Speed (cm/sec) | slope = $0.1827 \pm 0.04$ SE, $F(1,103) = 24.67$, $p < 0.0001$, $R^2 = 0.19$ | NA |
| 5G | Linear regression | Number of interactions | Time spent in centre (sec) | slope = $-0.1312 \pm 0.03$ SE, $F(1,103) = 16.17$, $p = 0.0001$, $R^2 = 0.14$ | NA |
| 5H | Linear regression | Duration of interactions (sec) | Time spent in centre (sec) | slope = $-0.4703 \pm 0.23$ SE, $F(1,103) = 4.069$, $p = 0.0463$, $R^2 = 0.04$ | NA |
| 5I | Linear regression | Inter-individual distance (cm) | Time spent in centre (sec) | slope = $-0.0033 \pm 0.0003$ SE, $F(1,103) = 78.52$, $p < 0.0001$, $R^2 = 0.43$ | NA |
| 5J | Linear regression | Nearest neighbour distance (cm) | Time spent in centre (sec) | slope = $-0.0001 \pm 0.0002$ SE,<br>$F(1,103) = 36.15$, $p < 0.0001$,<br>$R^2 = 0.26$ | NA |
| 6A | Linear regression | Communal egg-laying preference index | Egg laying latency (min) | slope = $0.0001 \pm 0.0001$ SE,<br>$F(1,103) = 1.172$, $p = 0.28$, $R^2 = 0.011$ | NA |
| 6B | Linear regression | Total number of interactions | Communal egg-laying preference index | slope = $4.337 \pm 8.965$ SE,<br>$F(1,103) = 0.234$, $p = 0.629$, $R^2 = 0.002$ | NA |
| 6C | Linear regression | Total number of interactions | Egg laying latency (min) | slope = $-0.021 \pm 0.013$ SE,<br>$F(1,103) = 2.424$, $p = 0.1225$,<br>$R^2 = 0.023$ | NA |

**Table S3:**
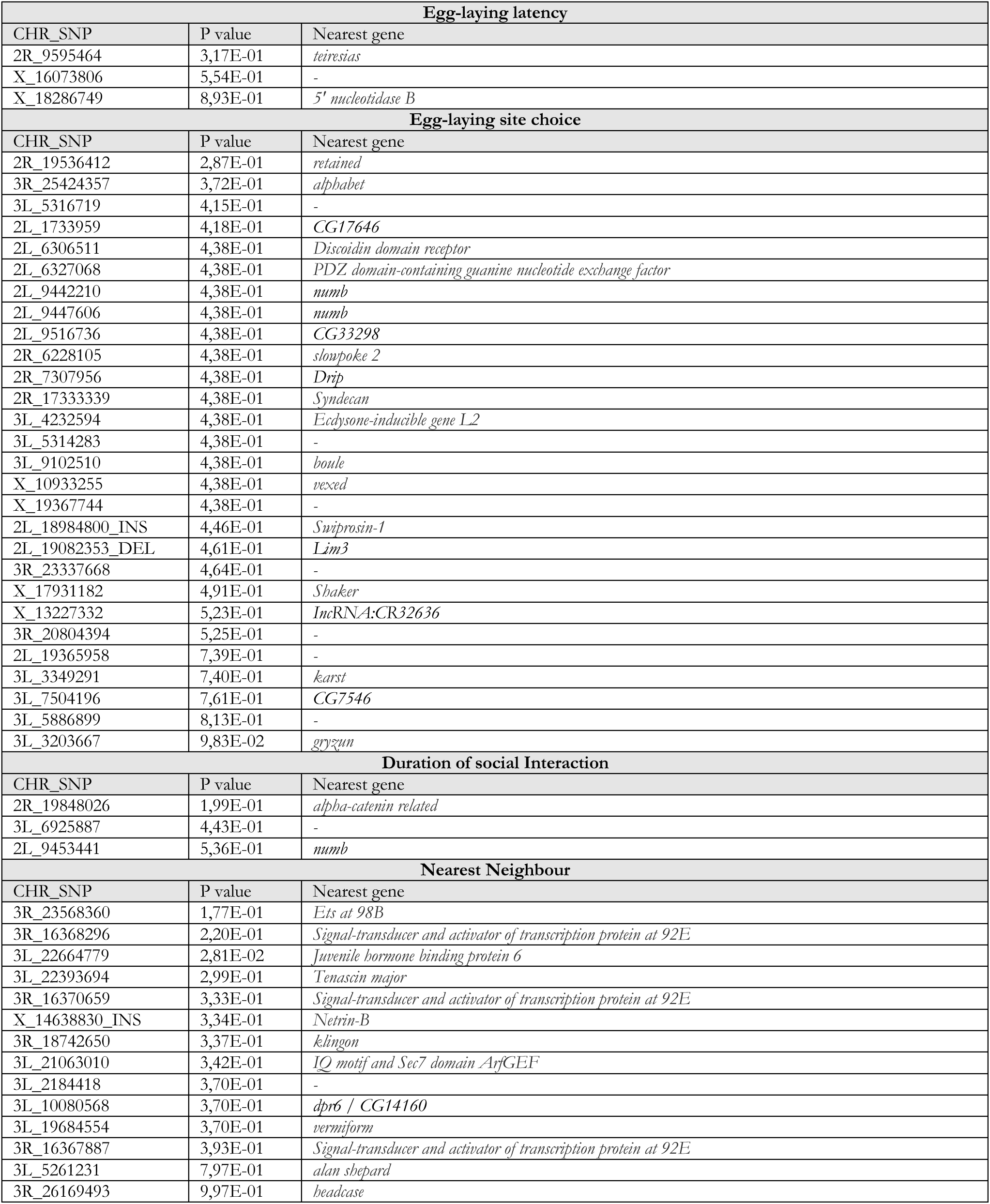
Significant Single Nucleotide polymorphisms (SNPs) from GWAS on the indicated phenotypes. Chromosomal location (CHR) is indicated. When followed by an “INS” indicates a nucleotide insertion and “DEL” indicates a deletion. P-value after Benjamini-Hochberg correction for false discovery rate is indicated, as well as the gene on which this SNP is found.

